# A Bait-and-Switch strategy links phenotypes to genes coding for Polymer-Degrading Enzymes in Intact Microbiomes

**DOI:** 10.1101/2025.10.09.681436

**Authors:** Colleen E. Yancey, Kyle D. Brumfield, Jackson Buss, Rita R. Colwell, Laurence Ettwiller

**Author notes:** Corresponding Authors: |.

## Abstract

Advances in next generation sequencing have made it possible to explore microbial community dynamics and regulation of functionally important genes through metagenomics and metatranscriptomics. However, the use of meta-omics to link enzyme function directly with complex, community-level phenotypes remain largely unexplored. To overcome this gap, we developed a novel framework that integrates ecological concepts by microbial community perturbation with association analysis to a targeted phenotype. Specifically, we introduce a hypothesis-free “bait and switch” strategy demonstrated through salt marsh soil microcosm pulse experiments to detect and characterize novel enzymes responsible for chitin degradation. Soil microbial communities were “baited” with shell compost, a chitin-rich substrate, to trigger community succession toward chitin degraders and gene upregulation of chitinases. A “switch” was then employed, by addition of glucose, inducing rapid downregulation of genes putatively responsible for chitin degradation. Results demonstrate the feasibility of this approach to identify functionally important enzymes, in this example, 48 hours after chitin addition. The bait and switch community perturbation provides a framework for discovery of polymer degrading enzymes present in complex microbial communities and serves as a proof of concept applicable for linking enzyme function with emergent community level phenotypes.

## Introduction

Complex polymers, both naturally occurring and artificially synthesized, comprise important substrates in natural systems, providing surfaces for biofilm formation as well as a source of critically important nutrient^1–4^. Cellulose, starch, lignin, and chitin derivatives are a few examples of natural polymers with broad environmental distribution. Chitin is the second most abundant polymer found in nature and is an important component in both carbon and nitrogen biogeochemical cycling. Despite its abundance, this polysaccharide does not accumulate naturally, suggesting its efficient turnover and degradation^5,6^. Microbial communities have evolved over time with such polymers common in their natural environments to produce enzymes that can degrade these substrates into oligomers and/or monomers that can be assimilated into the cell and metabolized.

Polymer degrading enzymes have significant application in material sciences, biotechnology, and remediation, depending on the polymer of interest^7,8^. However, accessing enzymatic diversity remains challenging, since conventional culture-based methods are estimated to recover *c.a* 1–2% of total environmental microorganisms under standard laboratory conditions^9,10^. Advances in molecular biology, metagenomics, metatranscriptomics, and computational approaches have greatly expanded our capacity to identify and characterize enzymes directly from environmental samples, bypassing the need for cultivation^11^. Nevertheless, these strategies remain largely dependent on existing enzyme databases, resulting in discoveries constrained to those enzymes with similarity to the already known, creating a “circular discovery loop”.

In contrast, microcosm pulse experiments offer a powerful approach to enzyme discovery by studying microbial communities under controlled, yet ecologically relevant conditions. These systems essentially preserve community structure while allowing targeted perturbation, such as nutrient amendment, to drive compositional and functional changes^12^. For example, crude oil addition in freshwater sediment microcosm experiments have been shown to stimulate microbial succession, selecting microbes containing hydrocarbon-degrading enzymes^13^. Such experiments highlight how dynamic microbial response can potentially be used for novel protein discovery without the need for isolation and cultivation. However, due to the complexity of microbial community composition, determining the direct link between a community-level shift and production of specific enzymes remains challenging and relies heavily on similarity to known proteins to prioritize which targets to validate experimentally.

In this study, a workflow was developed to identify novel polymer-degrading enzymes produced by soil microcosms, by employing a bait-and-switch pulse strategy coupled with phenotypic association analysis. As proof of concept, we focused on chitin-degrading enzymes in salt marsh soils. The experimental design involved “baiting” a microcosm with shell compost, a rich source of chitin, followed by a rapid “switch” to an alternative carbon source. Metagenomic Genome–Phenome Association (MetaGPA)^14^ was used to link protein domains with phenotypic traits, measured through community succession dynamics and transcriptional responses to bait- and-switch perturbation. With this approach, we successfully identified 190 known and putative chitinase genes, of which 27 were expressed for activity screening and 17 (*c.a* 63%) experimentally validated. Importantly, this hypothesis-free framework also enables tracking of global microbial community responses to carbon amendment (e.g., chitin, glucose). These results demonstrate the feasibility of combining a bait-and-switch experimental design with phenotype association analyses to accelerate enzyme discovery directly from environmental microbial communities.

## Results

### Bait and Switch Strategy

Pulse perturbation, namely substrate addition intended to transiently alter community composition, was applied to salt marsh soil microcosms to facilitate identification of novel-chitin-degrading enzymes by direct genotype association. To achieve this, we made the hypothesis that the addition of shell compost (predominantly chitin) would select for microorganisms utilizing chitin as a carbon source. Previous studies have shown that chitin serves as an effective selective substrate for chitinolytic microbes^15^ and that chitin-selective enrichment strategies have been successfully employed to favor chitinolytic microorganisms^16^.

The initial enrichment step, the “bait”, was designed to shift the microbial community composition toward enrichment of chitin-degrading taxa, as well as organisms benefiting indirectly from chitin as a nutrient source. Under the induced selective pressure, an increase in relative abundance of genes encoding chitin-degrading enzymes would be expected along with increased likelihood of detecting a larger diversity of enzymes by high-throughput DNA sequencing. Additionally, transcriptional activation of these genes should occur, resulting in increased relative transcript levels via high-throughput RNA sequencing following chitin amendment.

Following the bait, a “switch” strategy is applied to the same microcosms. This involves the introduction of a more readily metabolized carbon source, namely glucose, to stimulate downregulation of genes involved in chitin-degrading pathways. The term “switch” refers to the anticipated shift in transcriptional activity from chitin utilization in response to an alternate carbon substrate. Because transcriptional responses would be expected to occur more rapidly than community composition change, the switch strategy required sampling within hours following glucose amendment to capture any immediate changes in gene expression.

All experiments were performed in biological duplicates with controls that were not exposed to the bait and/or switch. The switch control comprised addition of the equivalent volume of ultrapure water as a substitute for the glucose solution. The overall experimental design is illustrated in Figure 1A. Sampling was at 0, 1, 4, 8, 24 and 48 hours after the bait and 1 and 4 (49 and 52 hours, total elapsed time) hours after the switch. DNA and RNA were extracted and metagenomic and metatranscriptomic libraries generated for all samples and analyzed using Illumina sequencing.

**Figure 1:**
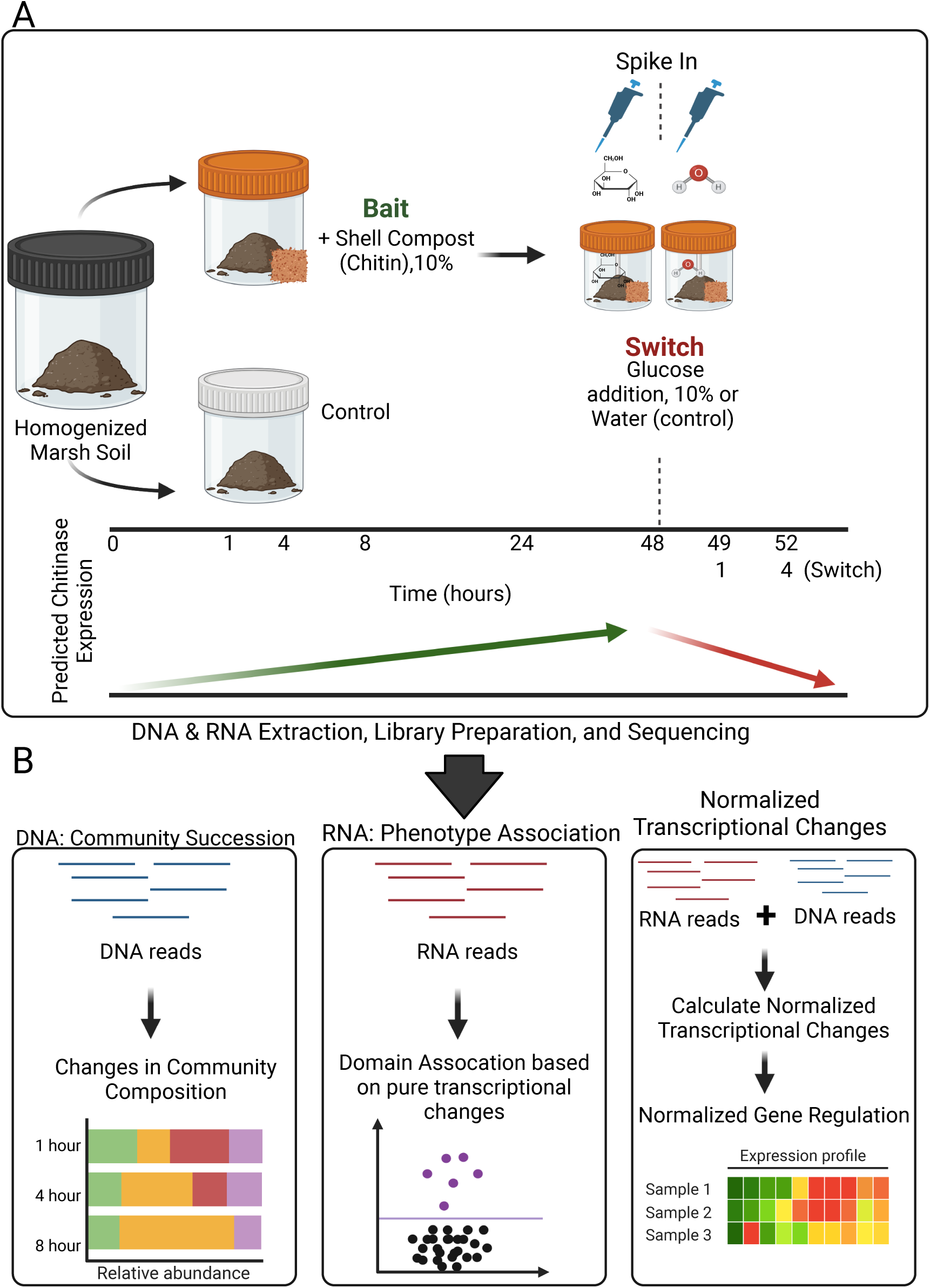
Overview of Experimental Design, Sampling Scheme, and Analysis Workflows. **A)** Microcosm pulse experiments were designed using the bait and switch model. At each designated timepoint in the sampling scheme, two grams of soil were collected in duplicate and processed for both RNA and DNA sequencing. **B)** Three association analyses were completed using the MetaGPA pipeline: DNA reads, RNA reads, and Normalized Transcriptional Changes (RNA:DNA ratios, see **Materials and Methods**).

### Chitin Bait Selectively Enriched for Chitin Degrading Taxa

To assess the effect of the bait and switch strategy on microbial community composition, alpha and beta diversity were calculated, employing read k-mer profiling of all metagenomic samples. Alpha diversity, or species diversity within a sample, was relatively the same with respect to genus for all samples within the first 8 hours. At 24 hours, alpha diversity decreased in chitin-treated samples (Fig. 2A), suggesting a selective enrichment of a limited subset of bacteria that possess the enzymatic machinery required for chitin degradation.

**Figure 2:**
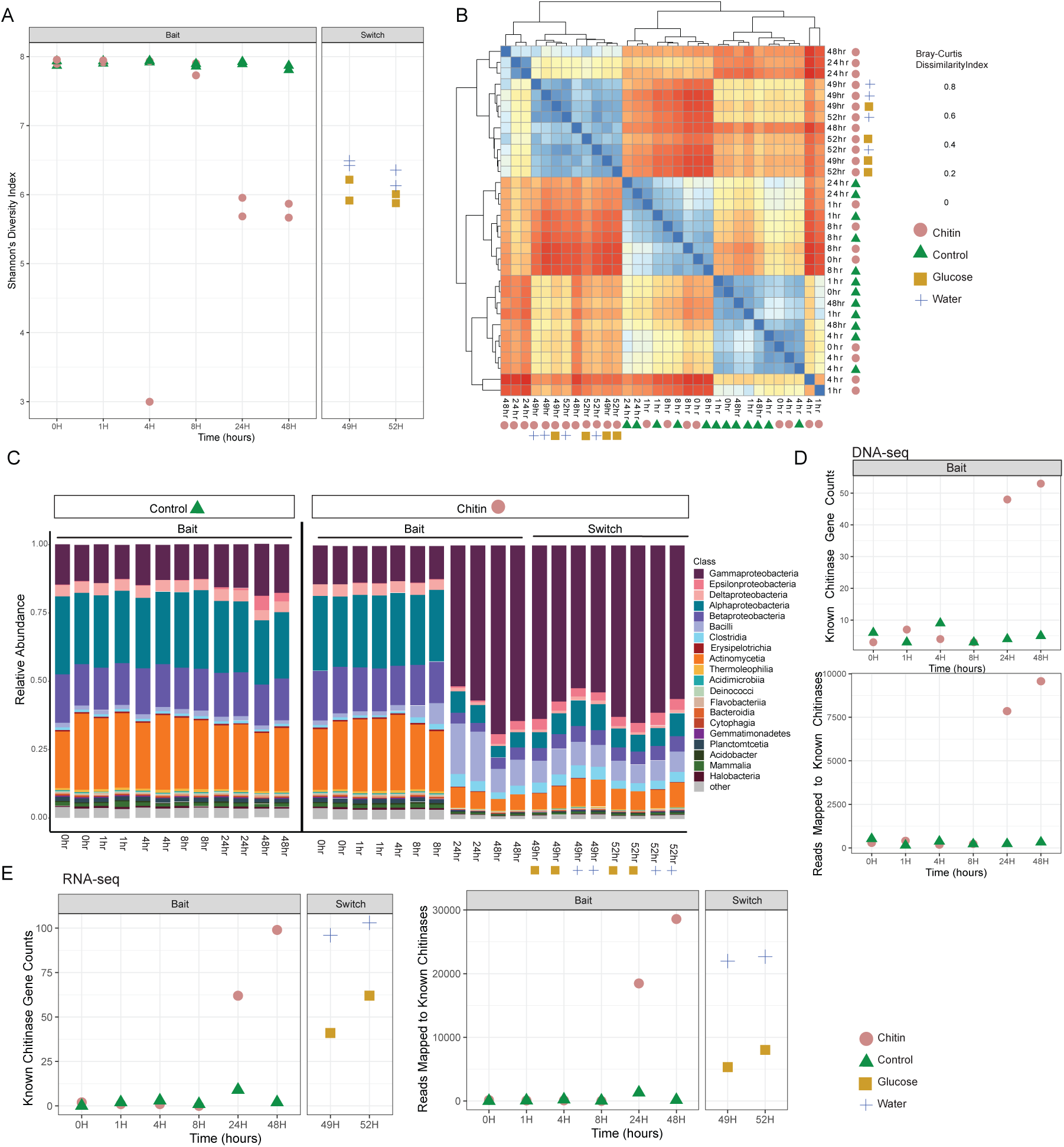
Soil Microbiome Community Changes during the experimental time course. **A)** Alpha Diversity, as measured by Shannon’s Diversity Index at the Genus level. **B)** Beta Diversity, as measured by Bray Curtis Dissimilarity at the Genus level. **C)** Relative abundance of different Classes. Diversity and community composition were calculated via k-mer frequency profiling using kraken2. **D)** Number (left) and total coverage (right) of homologs matching known Chitinase genes (from 30042170, Table 1) at different times of the bait experiment using DNA-seq data. **E)** same as **D)** but based on RNA-seq data.

**Table 1:**
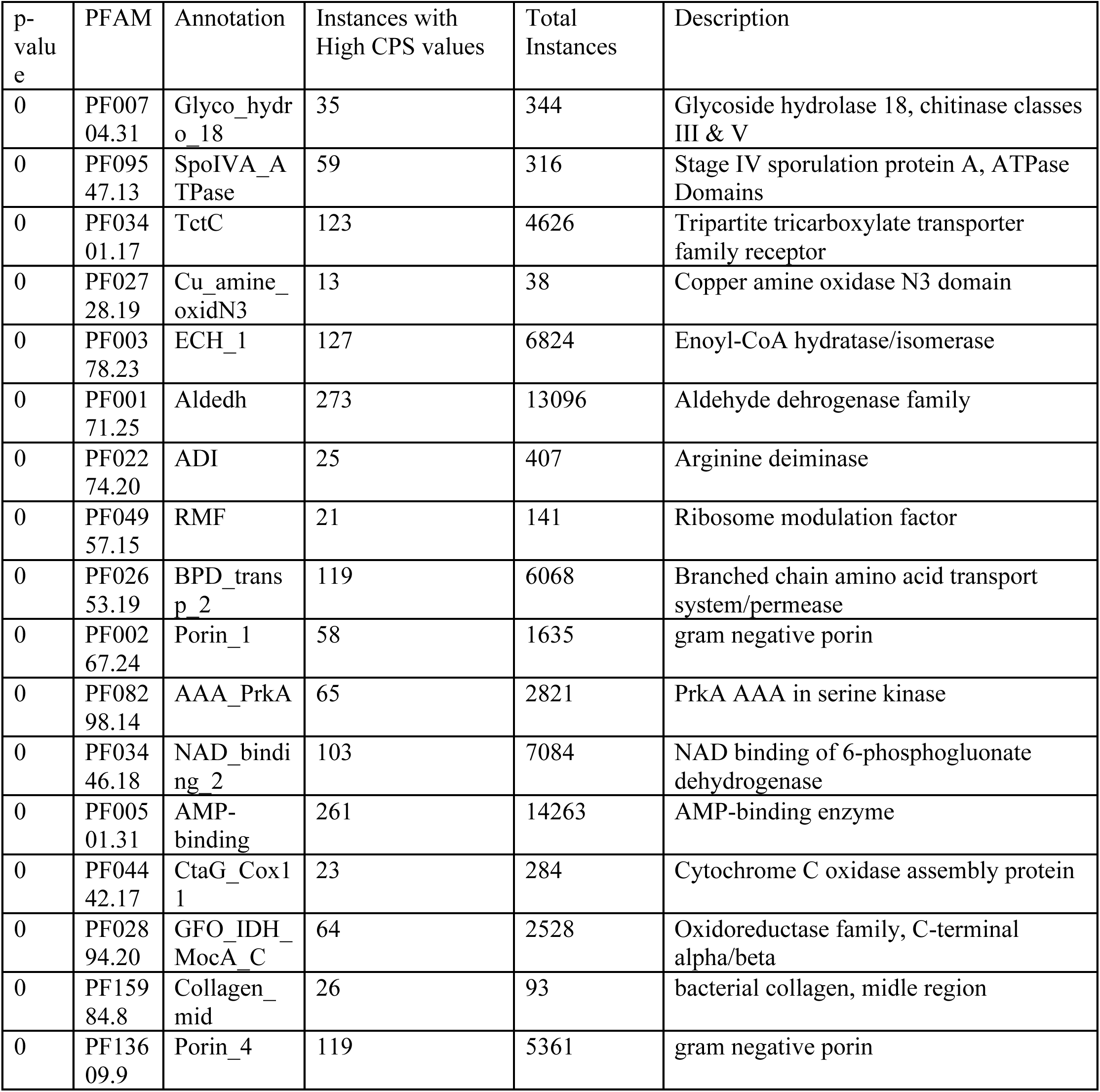
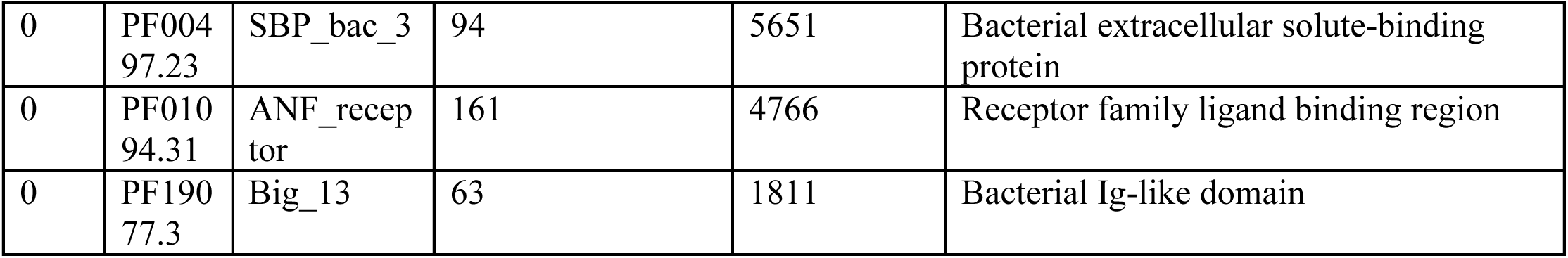
Top 20 Associated PFAM domains from the Combined Phenotype Association.

Three major groups, based on beta diversity, were determined by Bray-Curtis Dissimilarity analysis (Fig. 2B). Groups 2 and 3 include control and treated samples collected within the first 8 hours. Group 1 contained chitin-treated samples from time points of 24 hours or later, corresponding to decreased alpha diversity (Fig. 2B). At 24 hours and later, *Gammaproteobacteria* and *Bacilli* increased compared to control samples (Fig. 2C). Community composition did not change significantly following the glucose addition, as the short sampling interval (1–4 hours post-addition) was insufficient to capture detectable microbial succession (Fig. 2B, 2C).

### Genes coding for chitin degradation detected after chitin addition

A representative set of 15 experimentally validated chitinases^17^ was used to identify known chitinase coding genes via homology search. The number of genes homologous to validated chitinases increased 24 and 48 hours after chitin addition (Fig. 2D). Also, chitin amended samples exhibited a 10-fold increase in the number of genes compared to the original sample and a 17-fold increase relative to untreated controls. The relative abundance of genes, estimated by read mapping, rose 24 hours after chitin addition. By 48 hours, mapped reads were approximately 30-fold higher than the initial sample and untreated controls. (Fig. 2D). It is concluded that chitin addition during the bait period led to an increased abundance of genes predicted to encode chitin-degrading enzymes.

Similar patterns were observed with RNA-seq data, representing relative changes in transcription. Both the number of transcripts homologous to validated chitinases and total number of reads mapped, increased 24 and 48 hours after chitin addition. Relative transcript abundance of known chitinases decreased 1-4 hours after glucose addition. Addition of water did not alter transcript abundance of chitinase coding genes (Fig. 2E). These results support our hypothesis that chitin addition upregulates chitinase genes, and glucose addition stimulates rapid downregulation. Taken together, these results indicate both a selection for chitin degrading organisms and an apparent upregulation of transcripts coding for chitinolytic enzymes during the bait and a sharp downregulation of those transcripts during the switch. These findings support the use of phenotype–genotype association analyses to identify known and new chitinase genes. The phenotype of interest was a combination of both transcriptional activation in response to chitin addition and downregulation upon glucose addition.

### Phenotypic Association

Combined Phenotype Scores (CPS) were used to rank candidate transcripts associated with chitin degradation and identify domains (Table 1), an approach essential to capture the results of the bait and switch simultaneously. Specifically, transcripts showing strong upregulation during the bait phase (48 hours) and rapid downregulation during the switch (1 hour) had high CPS values (see **Material and Methods**). Transcripts annotated using the Protein Families Database (PFAM) and association analysis (modified from metaGPA) identified significantly enriched protein families among high-CPS transcripts. Recovered *de novo*, was an unbiased list of 105 associated protein domains (p-value=0, Tables 1 and S3), with the top hit, a glycoside hydrolase domain from family 18 (PF00704, Glyco_hydro_18, GH18), shown to be widely distributed in hydrolytic enzymes with chitinase or endo-N-acetyl-beta-D-glucosaminidase (ENGase) activity as well as chitinase-like lectins (chi-lectins/proteins (CLPs)^5,18^. Other highly ranked domains included those involved in solute binding and transport, and metabolite biosynthesis and utilization. A complete list of ranked domains is provided in Supplemental Table S3. For further phenotypic association analyses, associated domains were filtered by annotation for those identified as glycoside hydrolases (GHs), a broad group of enzymes in all domains of life, that hydrolyze glycosidic bonds, the expected mechanism of chitin degradation^18^.

### Association Analysis (MetaGPA/TPA) reveal enrichment of chitinase gene families

Next, we analyzed the association profile of GH18 and other GH domains across time during the bait and switch and observed significant associations 24-48 hours after chitin addition (bait). Significantly enriched domains included GH18, GH19, and GH20 using both RNA and DNA data. (Fig. 3A, B). This observation suggests both 1) an increase in the abundance of microbes harboring genes containing these GH domains, and 2) transcriptional upregulation of these genes. During the switch phase, one hour after glucose addition, domains GH18, GH19, GH20 and others were significantly depleted (Fig. 3B). Significant association in GH domains using transcription levels normalized to corresponding genomic abundances, highlighted “true” transcriptional changes as opposed to changes in community composition. GH18 exhibited the highest enrichment score 48 hours after chitin addition. This domain was also significantly depleted one and four hours after glucose addition (Fig. 3C). These additional analyses strengthen the evidence associating GH18 with chitin degradation (Table 1, Fig. 3).

**Figure 3:**
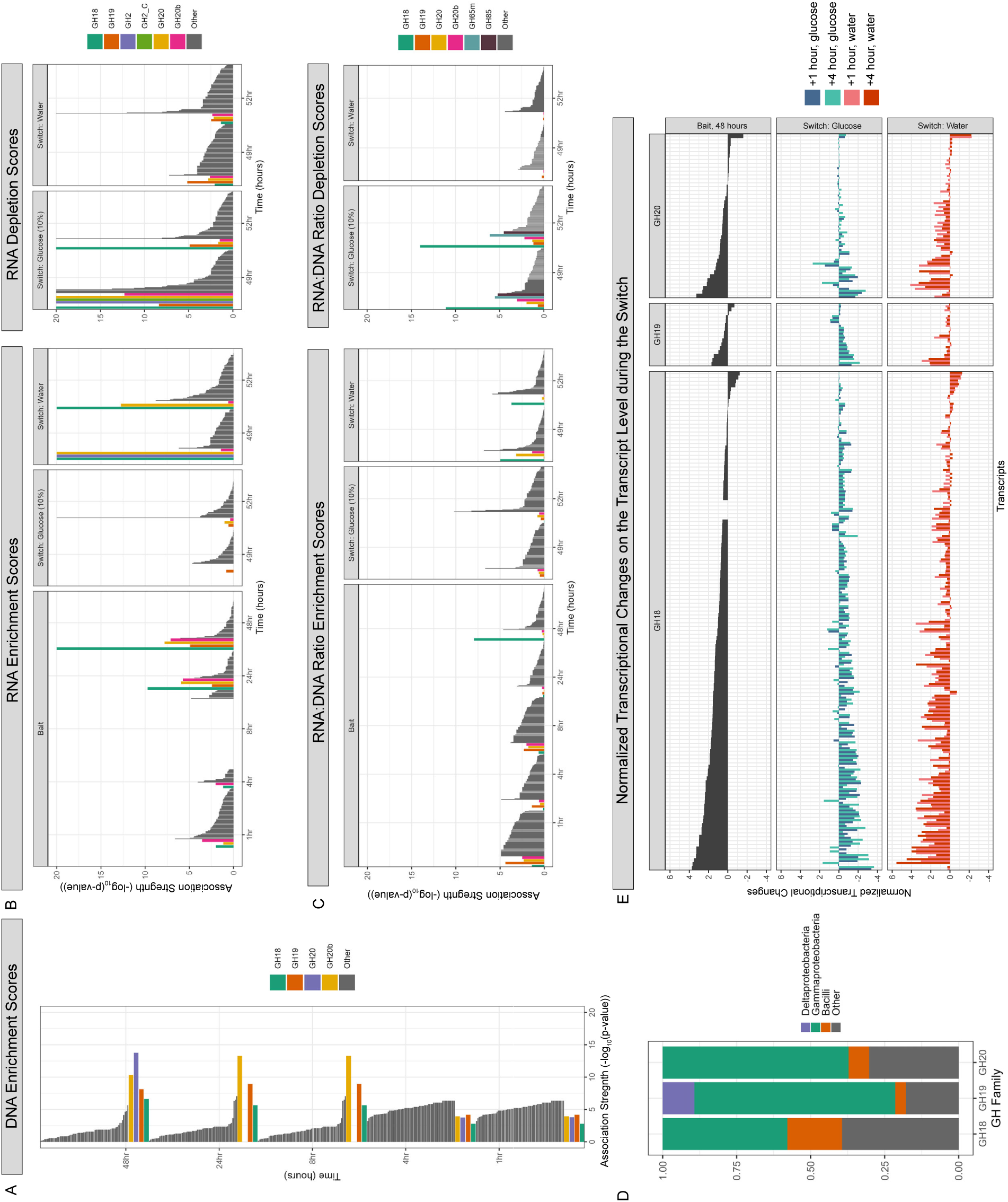
GH Association Scores across different analysis strategies. **A)** DNA enrichment scores during the bait phase of the experiment (1-48 hours post chitin enrichment). **B)** RNA association scores including enrichment scores during the bait phase (1-48 hours post chitin enrichment) as well as the switch phase (1-4 after glucose or water addition). Depletion scores for the switch phase are also shown. **C)** RNA:DNA Ratio, or normalized transcriptional changes, association scores. Enrichment scores during the bait and switch phase as well as depletion scores during the switch are shown. Domains are considered significantly associated if the association score (-log_10_(p-value) is equal or greater than 5. **D)** Relative abundance of identified transcripts assigned to bacterial Class for GH18, 19 and 20. **E)** Individual Normalized Transcriptional Changes for each transcript containing a GH18, GH19, or GH20 domain during the switch phase of the experiment. The top panel illustrates normalized transcriptional changes 48 hours after chitin addition. The middle and bottom panels illustrate normalized transcriptional changes between 48 hours after chitin addition and 1–4-hour post spike in (glucose or water).

### Regulation of Putative Chitinases Genes

Next, we identified all the genes that contained GH18, 19 or 20 domains. These genes were identified across several bacterial taxa, many of which are uncharacterized or uncultured (Fig. 3D). The majority of putative chitinase genes were upregulated 48 hours after chitin addition and downregulated one hour after glucose addition. (Fig. 3E, S1). Most candidate sequences contained a GH18 domain (EC 3.2.1.14, n=128), shown to have endochitinase, exochitinase, and N-acetylglucosaminidase activity^18^. The GH19 domain, described to have exochitinase activity^18^, was detected in fewer sequences (EC 3.2.1.14, n=19). Several sequences were found to contain GH20 domains (EC 3.2.1.52, n=43), which have N-acetylglucosaminadase activity, catalyzing breakdown of diacetylchitobiose in N-acetylglucosamine^18^. Most of the sequences were taxonomically assigned to *Gammaproteobacteria* and *Bacilli* (Fig. 3D), the relative abundance of which increased at 24-48 hours in samples treated with chitin (Fig. 2C).

### Experimental Validation of Putative Chitinases

Complete ORFs, containing GH18 domains, were expressed and screened for chitinolytic activity (see **Materials and Methods**). While these ORFs differ in sequence similarity and domain composition (Fig. S1, Fig. 4E), they exhibit the expected pattern of up- and downregulation in response to the bait and switch, respectively (Figure 3E). Screened sequences included bacterial Classes *Gammaproteobacteria* (11 sequences), *Bacilli* (6 sequences), *Unclassified* (6 sequences, unable to confidently assign taxonomy), *Bacteroidia* (2), and *Clostridia* (2). Of the 27 sequences screened, 17 exhibited chitinolytic activity. Proteins with greatest relative activity, especially exochitinase and endochitinase functionality, are from *Gammaproteobacteria*. In addition to containing GH18 domains, several of the proteins also contained chitinase A and carbohydrate binding module family 5/12 domains (ChiA: PF08329 and CBM: PF02839, Fig. 4E). Compared to the chitinase assay positive control, chitinases from *Gammaproteobacteria* had relatively higher exo- and endochitinase activity, but not β-N-acetylglucosaminidase activity. Relatively low β-N-acetylglucosaminidase activity was observed for GH18_19, from *Bacilli*, that lacked ChiA and CBM domains. Two proteins, from *Clostridia* (GH18_23, GH18_24), exhibited relatively high and very high exochitinase activity, despite lacking ChiA and CBM (Fig. 4). GH18_23 and GH18_24, homologous to *Clostridium* spp. CotE proteins, are important proteins in spore formation^19,20^.

**Figure 4:**
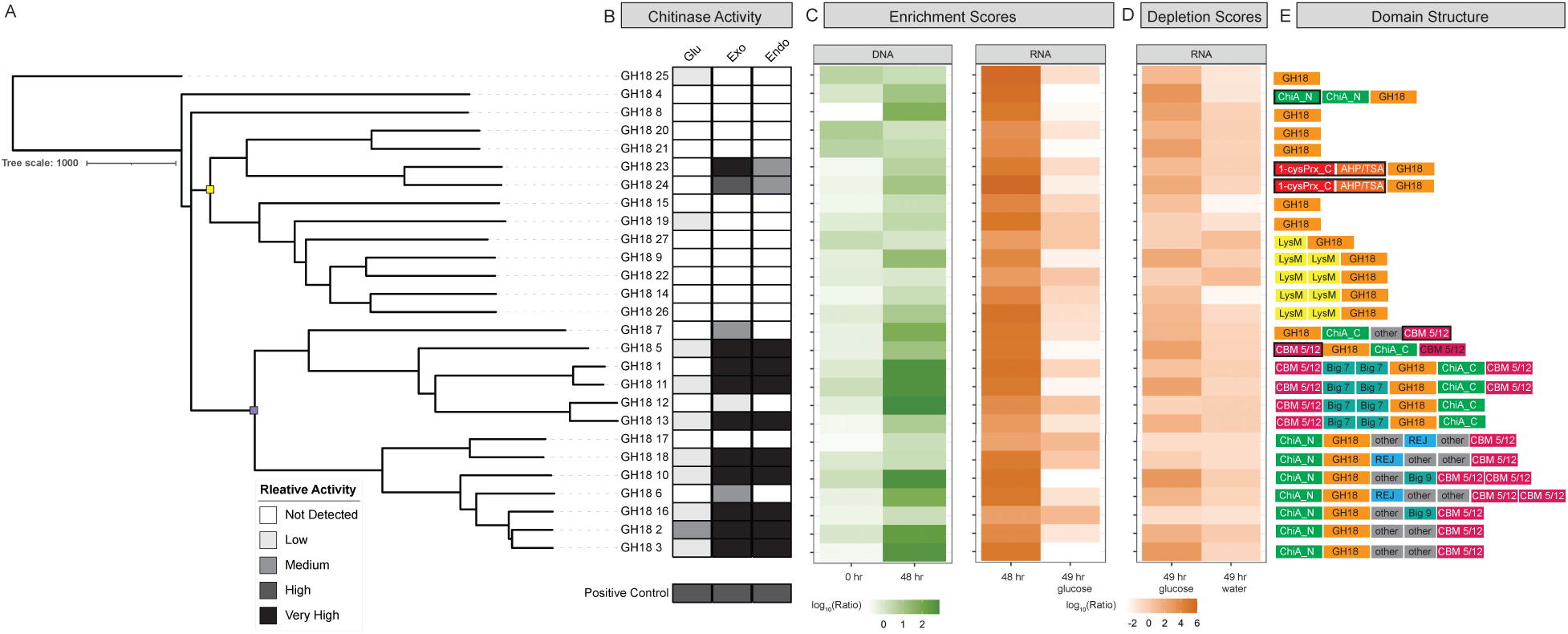
experimental validation of 27 novel chitinase candidates. **A)** A phylogenetic tree for the 27 GH18 domain-containing proteins is shown with relative chitinase activity. The phylogenetic tree was generated using a MAFFT alignment and neighbor joining tree algorithm (1000 iterations). The yellow and purple nodes contain sequences from bacteria from the classes Bacilli and Gammaproteobacteria, respectively. **B)** Relative chitinase activity for β-N-acetylglucoasmindase (Glu), exochitinase (Exo), and endochitinase (Endo) are depicted on the right. Relative activity was categorized based on the measured release of 4MU (ng): Not Detected <0 ; Low <0, ≤ 10; Medium: > 10, ≤ 100; High: >100, ≤500; Very High: > 500. **C)** Enrichment scores are shown for metaGPA analysis completed on DNA or RNA respectively. DNA enrichment scores are shown for the bait phase of the experiment: 0 hours and 48 hours post chitin addition. RNA enrichment scores are shown for the switch phase of the experiment: 48 hours after chitin addition and 1 hour after glucose addition. **D)** RNA depletion scores are shown for the switch phase of the experiment: 1 hour after glucose or water addition as compared to 48 hours post chitin addition. Schematic domain structure for each protein sequence is shown to the right of depletion scores. **E)** Domains include glycoside hydrolase 18 (GH18), Chitinase A, N terminal (ChiA_N), Chitinase A, C terminal (ChiA_C), LysM domain (LysM), carbohydrate binding module family 5/12 (CBM 5/12), bacterial iG-like domains (Big_7, Big_9), the REG domain (REG), 1-cys peroxiredoxin C-terminal (1-cysPrx), AhpC/TSA family (AhpC/TSA), and other (all other domains). Domains outlined in black boxes indicate domain arrangement that has not previously been seen before in experimentally validated chitinases.

## Discussion

Bait and switch pulse experiments were carried out with domain association analyses to establish a framework for polymer degrading enzyme discovery directly from intact microbial communities. The approach included three integrated experimental phases. First, shifts in community composition were measured by changes in relative DNA abundance, serving as a proxy for microbial succession in response to addition of the bait. Next, a gene expression baseline within the enriched community was established, enabling hypothesis free association of genotype to the desired phenotype (polymer degradation). Finally, a switch, or deliberate perturbation of the microbial environment induced rapid downregulation of associated genes. Together, these three phases: community succession, gene expression in response to the bait, and gene downregulation in response to the switch, provide a robust procedure to determine the association of microbial phenotypes with genes that encode targeted functions. Thereby, successful identification of a list of putative chitin-degrading sequences from salt marsh soils from both known and unclassified bacteria was established (Fig. 3,4). Notably, we observe an enrichment for *Gammaproteobacteria*, *Bacilli*, and other taxa, which are known to naturally occur in these systems, associated with chitin substrates^21^. Expression of sequences confirmed activity and validity of the approach to identify target enzymes of interest (Fig. 4).

The workflow outlined here allows discovery and characterization of novel enzymes that may have been overlooked by more established approaches. Chitinolytic activity was confirmed for 17 of 27 constructs screened. Of the 17, five sequences had >98% homology with previously validated chitinases and the other 12 contained glycoside hydrolase 18 domains. Identification of associated proteins from unculturable or poorly characterized microbial taxa can also be observed. For example, six of the 27 GH18-domain proteins tested for activity lacked formal taxonomic assignment, and two of which (GH18_5 and GH18_7), exhibited measurable chitinolytic activity (Fig. 4).

Finally, proteins with novel domain arrangement, compared to experimentally validated chitinases, can be identified. Additional carbohydrate-binding domains flanking the GH18 domain, not previously reported in characterized chitinases, retain chitinolytic activity, as observed in GH18_5 and GH18_1 (Fig. 4E). These additional domains may enhance activity via higher affinity for substrate binding^22^. Chitinolytic activity was also observed for GH18_23 and GH18_24 which have homology to *Clostridium* spp. CotE proteins. This protein has been previously described as a bifunctional peroxiredoxin-chitinase^20^, with possible implication in spore binding to mucus during infection^23^. While recombinant mutants of this protein containing only the GH18 domain have been previously demonstrated to be chitinolytic^19^, to our knowledge, this is the first demonstration of chitinolytic activity of the complete protein sequence containing both the peroxiredoxin and chitinase domains (Fig. 4).

The results presented have shown that it is possible to implement microbial ecology and phenotype associations to identify functionally important enzymes of interest. While chitin degradation was used to develop the workflow, the approach can be used for the discovery of enzymes degrading other polymers, e.g. plastics^24^. The three phase approach of the bait and switch is even more critical when functional characteristics of domains driving observed phenotypes are not constrained, in contrast to chitin-degrading enzymes. Additionally, relying on association at the *domain* level, rather than gene databases, provides greater resolution and targeting for the discovery of novel enzymes and functionalities. The results of this study emphasize the importance of using paired multi-omic data as community succession events can be assessed through metagenomic analysis, but changes in gene regulation, especially after the switch, are rapid (1-4 hours) and must be evaluated via metranscriptomics (Fig. 3). This approach has the potential to rapidly improve enzyme discovery and characterization timelines. It is possible to screen an increased number of enzymes from uncharacterized taxa, which are believed to make up the majority of microbial diversity^25–27^, thereby releasing bottlenecks introduced by challenges in cultivation^9,10^. Given the multifaceted layers for discovery and characterization within this singular, hypothesis-free pipeline, its expansion and application provide the potential for continued enzyme discovery.

## Methods

### Sample Site

Soils were collected from the Great Marsh, Ipswich MA, USA (42.6672° N, 70.8111° W), which spans much of the North Shore of Massachusetts. The sample area consists of an open marsh drained by the Labor-in-Vain Creek and experiences flooding during high tide and drainage during low tide. The site is largely dominated by high marsh (greater than 1.3m) and experiences semidiurnal tides with an average mean range of 2.5-2.9m^28,29^. The marsh is dominated by the salt meadow cordgrass, *Sporobolus pumilus*^29,30^, and sediments are carbon-rich compared to other salt marshes in the eastern United States^29^.

Sampling was completed on 2, October 2023, at 7:15 am, during low tide. During the time of sampling, the pH of the soil was about 6, as approximated with VWR pH-Test strips (BDH35309.606). Approximately 2 kg of soil was collected at 0-10cm depth, adjacent to *S. pumilus* growth, which provides a suitable habitat for chitin-rich organisms such as crustaceans and mollusks^31^. Samples were stored in sterile high-density polypropylene (HDPE) bottles during transport.

### Microcosm Experiments

Collected soil was homogenized and sieved through a 2.36 mm sieve. Stones, sticks, root nodules, and other litter were removed. Two microcosms were set up, containing 600g of homogenized soil, each in sterile glass containers. The first microcosm served as a control. The second microcosm was amended with 10% w/w sterile, ground, Neptune’s Harvest Crab and Lobster Shell Compost (Gloucester, MA, USA) which served as the chitin amendment (treatment). Shell compost was selected versus purified or synthetically derived chitin to simulate chitin substrates found naturally in the environment. It has been shown that purified and chemically derived chitin is less preferred by microbes for metabolism and degradation^32,33^. Both microcosms were mixed thoroughly with sterile spatulas to ensure homogenization after addition. The experimental design and sampling scheme are illustrated in Figure 1.

For sample collection, microcosms were mixed with sterile spatulas and destructively sampled at specified time points (0, 1, 4, 8, 24, 48, 59, and 52 hours). At each time point, two-gram aliquots were collected in duplicate, flash frozen in dry ice-ethanol slurry, and stored at - 80°C until further analysis. At 48 hours, another round of sampling was completed prior to glucose or water addition. The remaining soil in both the control and chitin amended microcosms were split into two, new, sterile containers each containing half of the original soil (∼250 g), thereby providing two microcosms with the control soils, and two microcosms with the chitin amended soils. For one of each microcosm group (treatment, or control), a sterile 50% w/v glucose solution was added to obtain 10% w/w glucose. For the remaining microcosm groups (one treatment, one control), the same volume of sterile, deionized water was added to control for soil rehydration. All four microcosms were thoroughly mixed with sterile spatulas, and the sampling scheme continued until the completion of the experiment.

### Library Preparation

RNA for metatranscriptomic sequencing and analysis was extracted and prepared as described. RNeasy® PowerSoil® Total RNA kits (12866, Qiagen, Hilden, Germany) were used to extract total RNA from 2 grams of soil for each replicate according to the manufacturer’s instructions, with some modifications. The lysis/bead bashing step was increased to 20 minutes. During the isopropanol precipitation, samples were incubated at -20°C for 45 minutes. To remove any residual polyphenolic or humic acids that would interfere with downstream library preparation, the Zymo OneStep PCR Inhibitor Removal Kit (D6030, Zymo Research Corp, Irvine, CA, USA) was used to treat and concentrate purified RNA according to the manufacturer’s instructions. RNA concentration and purity ratios were determined using a NanoDrop One (ThermoScientific, Waltham, MA, USA). Samples were stored at -80°C until library preparation was completed. For each sample, around 800 ng of total RNA was used for library preparation. Ribosomal depletion was completed using the NEBNext® rRNA Depletion Kit for Bacteria (E7850, New England Biolabs, Ipswich, MA, USA) according to the manufacturer’s instructions. Libraries with 200 bp inserts were prepared with the NEBNext® Ultra II Directional RNA Library Prep Kit (E7765, New England Biolabs, Ipswich, MA, USA). NEBNext® Multiplex Oligos for Illumina® (E7500, New England Biolabs, Ipswich, MA, USA) were used for barcoding and PCR enrichment. Library concentration, size, and quality were assessed using a TapeStation 4200 with a D1000 High Sensitivity Screen Tape (5067-5584 Agilent, Lexington, MA, USA).

DNA for paired metagenomic analyses was also extracted and prepared for sequencing. DNeasy® PowerSoil® Pro Kits (47016, Qiagen, Hilden, Germany) were used to extract total DNA from 0.25 g of soil per replicate according to the manufacturer’s instructions. As with RNA preparation, OneStep PCR Inhibitor Removal Kits (D6030, Zymo Research Corp, Irvine, CA, USA) were used to treat and concentrate DNA samples according to manufacturer instructions. DNA concentration and purity ratios were determined using a NanoDrop One (ThermoScientific, Waltham, MA, USA). Samples were stored at -20°C until further library preparation was completed. For each sample preparation, 50 ng of purified DNA was sheared to generate 200 bp inserts using a Covaris SR Focused Ultrasonicator (Covaris, Woburn, MA, USA). Libraries were generated using the NEBNext® Ultra™ II DNA Library Prep Kit for Illumina® (E7645, New England Biolabs, Ipswich, MA, USA), according to manufacturer instructions. NEBNext® Multiplex Oligos for Illumina® (E7500, New England Biolabs, Ipswich, MA, USA) were used for barcoding and PCR enrichment. Library concentration, size, and quality were assessed using a TapeStation 4200 with a D1000 High Sensitivity Screen Tape (5067-5584 Agilent, Lexington, MA, USA). For both DNA and RNA, libraries were sequenced at the New England Biolabs Sequencing Core, PE 2X100 on the NextSeq or NovaSeq S2/SP targeting 50 and 100 million reads, respectively (Illumina, San Diego, CA, USA).

### Bioinformatic Analysis

Relative changes in community composition in response to treatment were determined via metagenomic k-mer profiling using Kraken2 under default settings, using the Standard Collection database built for 100-mers (accessed October 2024)^34^. Alpha diversity as measured using Shannon’s Diversity Index, and Beta Diversity, as measured by Bray-Curtis Dissimilarity, were also calculated for each metagenomic sample using KrakenTools^34^. Calculations were completed at the genus level using k-mer profiles and KrakenTools to assess shifts in community composition. To assess the frequency of known chitinase gene enrichment or expression in response to treatment, the following analyses were completed. Reads from replicates were combined and downsampled to 30 million reads using seqtk sample (v1.4-r122). Assembly was performed with metaSPAdes (v3.15.3)^35^. Representative chitinases with experimentally validated activity were retrieved from a previous study^17^, and corresponding amino acid sequences were obtained from the Protein Data Bank (PDB) database (accession numbers: 1E15, 1WVU, 2CJL, 2Z37, 3CQL, 3HBD, 3G6L, 3IWR, 3N11, 3ALF, 3AQU, 3OA5, 3SIM, 2Y8V, 4AXN). Homologs of these chitinases were identified in the microcosm metagenomic and metatranscriptomic datasets from both chitin-treated and control samples at each time point using tblastn (BLAST+ v2.16.0, default parameters). Hits with *e*-values < 0.001 were combined and merged using the bedtools merge function (v2.27.1). For each dataset, the total number of hits and the corresponding read coverage were recorded.

Phenotype association analyses were completed using the core MetaGPA pipeline as previously described^14^ on metagenomes (DNA), metatranscriptomes (RNA), and normalized transcriptional changes (RNA:DNA ratios). Analyses of each dataset were completed to determine whether association at the gene level (representing changes in community composition), transcript level (representing changes in gene expression), or normalized transcriptional changes (representing changes in both community composition and gene expression) would provide the strongest signal for domain associations (Fig. 1B). The MetaGPA pipeline can be accessed at https://github.com/ge-a/metaGPA2.

Phenotype association analyses using MetaGPA were completed with the following modifications. For both metagenomic and metranscriptomic samples, technical duplicates were used to generate assemblies for treatment and control samples. For each time point, reads were assembled using metaSPADES (metagenomes) or directional rnaSPADES (metatransciptomes) under default settings (v.3.15.3)^35,36^. Assemblies from the treatment and control were combined and dereplicated at 99% identity and 90% alignment length using cd-hit (v4.8.1)^37^ to generate a non-redundant list of contiguous sequences (contigs) or transcripts observed in either the control or treatment, or both. Replicate read files from the control and treatment were individually mapped to the non-redundant, combined assembly using bwa-aln (v0.7.15)^38^. Read counts for each replicate per experimental group were averaged and used to calculate ratios between the treatment and control to determine if contigs were enriched or depleted in treatment samples. If the ratio of reads mapped between the treatment and control were greater than 3.0, the contig was considered enriched. Each contig was annotated using the Protein Family Databases (PFAM, v35.0), for protein domains^39^ using Hidden Markov Modelling (HMM) via hmmer (v3.3.2). To ensure accurate annotations, PFAM assignments were kept for further analysis if the annotation e-value score was ≤ 1.00 e-05. Fisher’s exact test was used to determine if the number of contigs observed in the treatment versus control was statistically significant. If contigs or transcripts were significantly abundant or depleted in the treatment, they were considered enriched or depleted in the chitin amended soils, respectively.

To assess whether significant associations observed in the classic MetaGPA workflow were a result of pure change in transcript abundance, community succession selecting for chitin degraders, or both, additional analyses were completed. First, both metagenomic and metatranscriptomic reads were mapped to assembled transcripts respective of sampling time. Each sample was normalized for sequencing depth, and a ratio of normalized RNA to DNA was generated for each contig. This ratio represents relative expression effort, correcting for differences in community composition across samples and variation in sequencing depth.

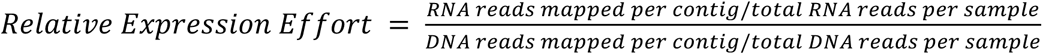

Ratios of relative expression effort between treatment and control samples were calculated to determine which genes were upregulated or downregulated in treatment samples compared to control. These normalized transcriptional changes were used to recalculate enrichment scores and determine which domains were differentially associated across different samples.

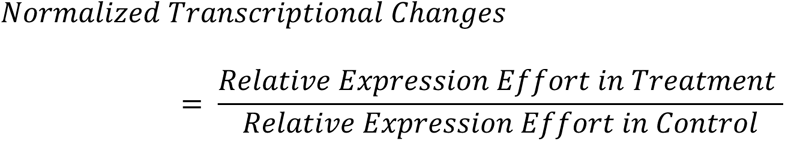

### Identifying Associated Candidate Domains for Screening

To integrate information from both the bait and switch to identify candidate chitinolytic domains, a scored phenotype association method was used. The scored phenotypes were applied to domain association results using metatranscriptomes from 48 hours after chitin addition (bait, Table S1) and 1 hour after glucose addition (switch, Table S2). If the ratio during the bait and switch was 3 and 0.33 respectively, a score of 10 was assigned indicating that both an upregulation during the bait and downregulation during the switch. If only one or neither condition was met, a score of 0 was assigned. These scores were used to complete domain association once more with a cutoff of 5, resulting in a list of domains associated with desired phenotypes in both the bait and the switch. (Table 1, S3).

Associated domains were further filtered based on annotation to identify candidate chitinase sequences. Results were filtered and domains that were annotated as glycoside hydrolases (GHs) were kept for further analysis. Open reading frames from transcripts from the 48-hour assembly that contained domains of interest were additionally used to build sequence similarity networks using the EFI Enzyme Similarity Tool v.2025_03^40,41^. Edges were drawn between nodes with an alignment score threshold of at least 35%, and networks were visualized using Cytoscape v. 3.10.3^42^.

### Expression and Activity of Purified Chitinases

To validate putatively identified chitinases, 27 predicted protein sequences were screened for activity. These sequences were selected based on the following criteria: sequences needed to be complete open reading frames (ORF) and annotated to contain a glycoside hydrolase 18 (GH18) domain. To determine if these identified sequences exhibited novel domain arrangements or structure, AlphaFold2 v.2.3.0 models were built for each predicted protein, and the Protein Databank (PDB) (accessed August 2025) queried with these models using FoldSeek v.10 under default parameters. Additionally, sequences were also aligned to the 444 experimentally characterized protein sequences containing GH18 domains found on the Cazy database (accessed 2 September 2025).

Gene blocks containing candidate chitinase sequences were assembled into the pet28a(+) backbone using the NEBuilder Hifi DNA Assembly Kit (E2621, New England Biolabs, Ipswich, MA, USA). Candidate chitinases were then synthesized *in vitro*. Approximately 1 µg of plasmid DNA was added to an NEBExpress Cell-free *E. coli* Protein Synthesis System reaction (E5360, New England Biolabs, Ipswich, MA, USA), scaled for a total reaction volume of 100 µL. Each reaction was supplemented with PureExpress Disulfide Bond Enhancer (E6820, New England Biolabs, Ipswich, MA, USA) to ensure proper folding of chitinase proteins, which contain disulfide bonds. The IVTT reaction was incubated at 25°C, with vigorous shaking for 24 hours. Expressed candidate chitinases were purified via his-tag affinity using NEBExpress Ni-NTA Magnetic Beads (S1423, New England Biolabs, Ipswich, MA, USA). Proteins were eluted with elution buffer consisting of 20 mM sodium phosphate, 300 mM NaCl, 300 mM imidazole, pH 7.4. Protein expression was visualized via SDS-PAGE (Fig. S2).

Purified proteins were screened for chitinase activity using the Fluorometric Chitinase Assay Kit (CS1030, Sigma Aldrich, St. Louis, MO, USA) according to the manufacturer’s instructions. The assay relies on enzymatic hydrolysis to release 4-methylumbelliferone (4MU) from labelled substrates to screen for exochitinase (4-Methylumbelliferyl N,N′-diacetyl-β-D-chitobioside), endochitinase (4-Methylumbelliferyl β-D-N,N′,N′′-triacetylchitotriose) and β-N-acetylglucosaminidase (4-Methylumbelliferyl N-acetyl-β-D-glucosaminide) activity. All three substrates were screened for each protein to fully characterize chitinolytic activity of the candidate proteins. In reference to the assay positive control chitinase from *Trichoderma viride* (C6242, Sigma Aldrich, St. Louis, MO, USA), relative levels of chitinolytic activity were determined for each protein sequence and substrate screened. Measurements of 4MU (ng) released below 0 were designated as not detected. The following designations were assigned based on measured release of 4MU – low: greater than 0 ng and less than or equal to 10 ng, medium: greater than 10 ng and less than or equal to 100 ng, high: greater than 100 ng and less than or equal to 500 ng, and very high: greater than 500 ng. The assay control chitinase exhibited high activity for each substrate tested.

## Acknowledgements

We are grateful to the following for providing materials, services, and support during this project. We thank the NEB Sequencing Core and NEB IT Department for their assistance in sequencing and technical support. We thank Paula Magnelli for critical suggestions and support. We thank Jennifer Ong, Katherine O’Toole, and Rebekah Silva who provided suggestions for protein expression. We are also grateful to Andy Ge for computational and bioinformatic support.

## Data Availability

Raw reads are available on the NCBI Sequencing Read Archive (SRA) under BioProject: PRJNA1293438.

---

**Figure S1:**
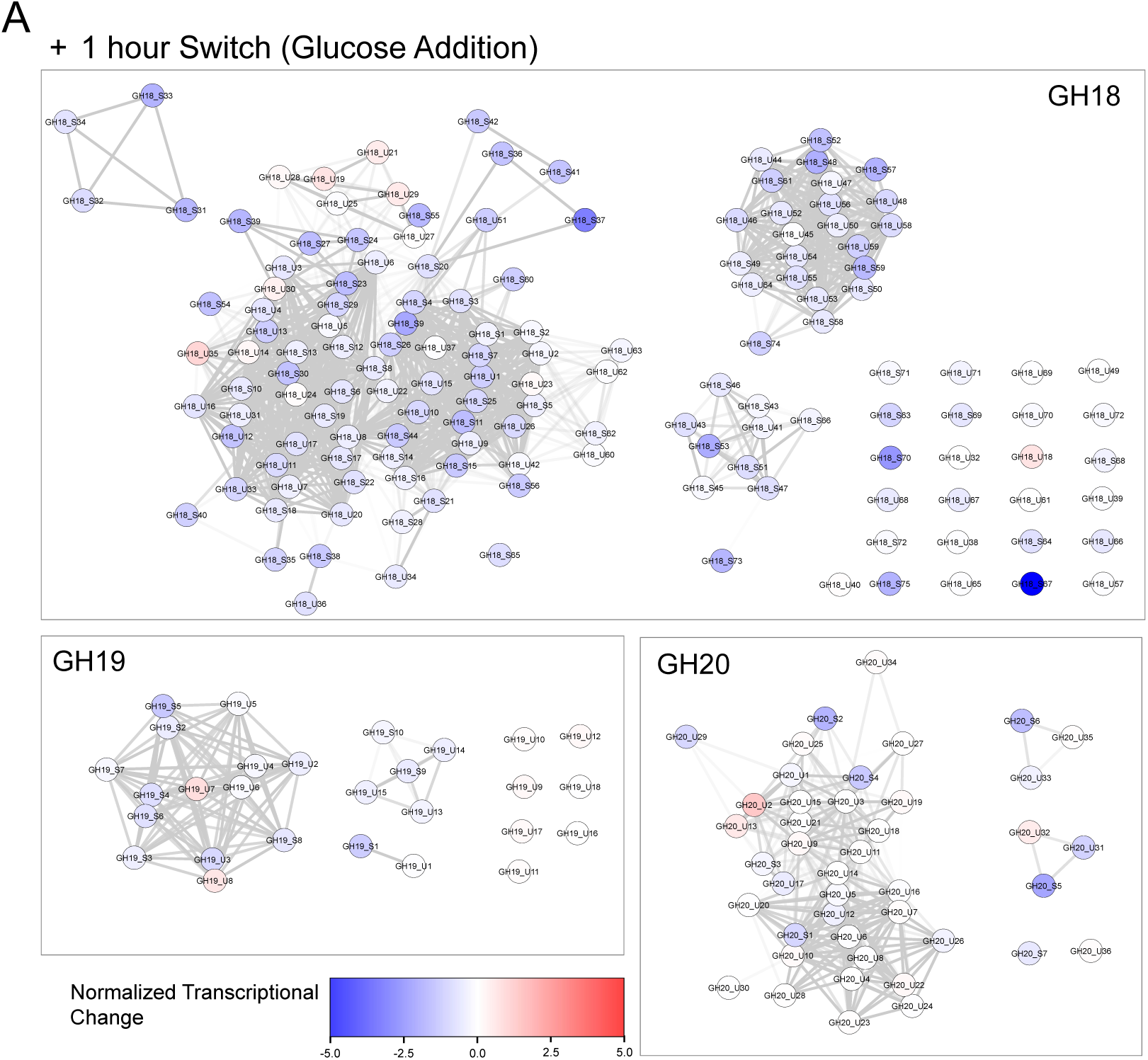
Similarity networks of putatively Identified Chitinases. A) Sequence similarity network for amino acid sequence predicted to contain GH18, GH19, and GH20 domains. Each node is colored by normalized transcriptional changes 1 hour after glucose addition (switch). Sequence similarity between proteins is represented by edge thickness.

**Figure S2:**
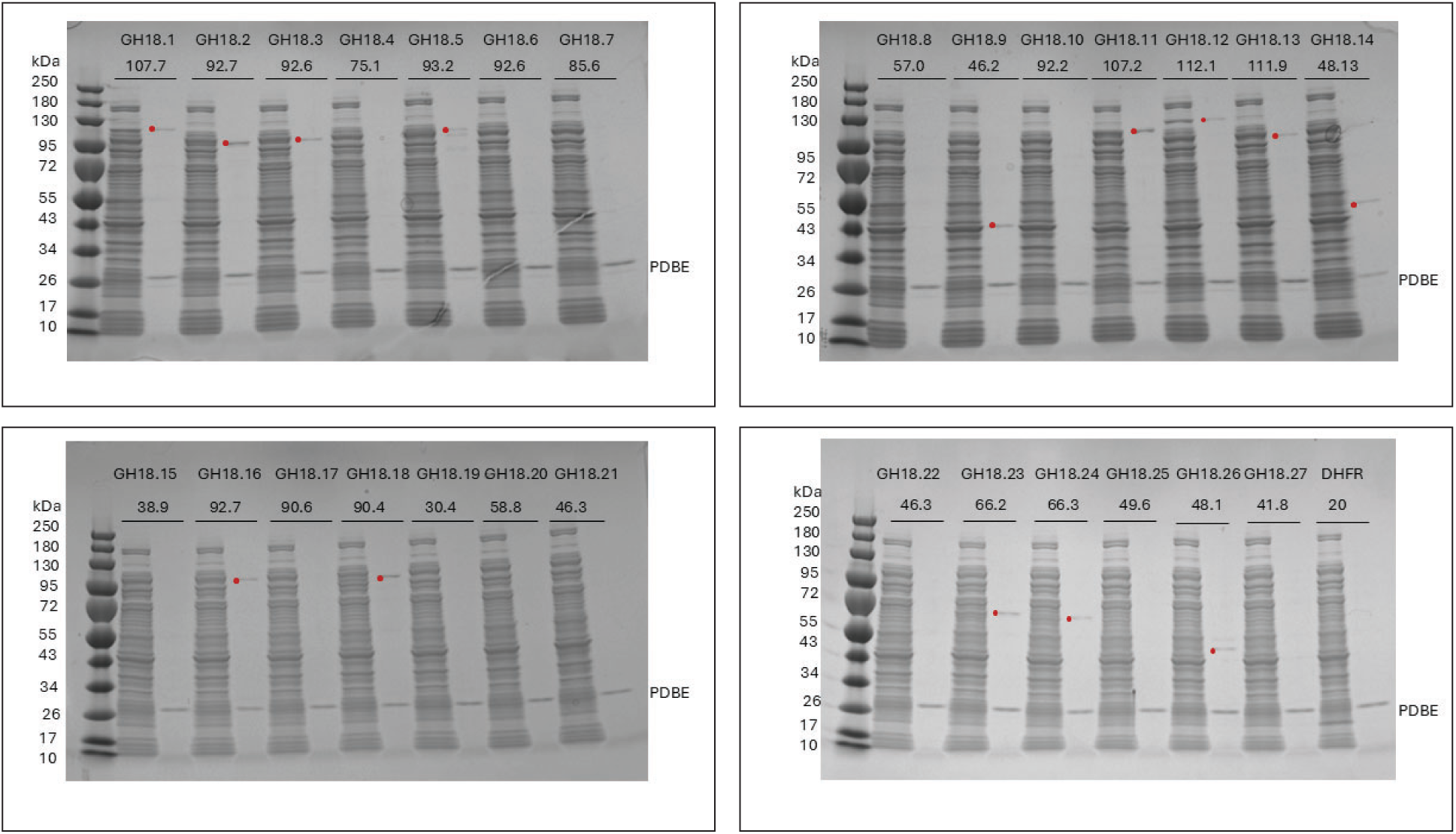
SDS-PAGE Gels of expressed chitinase candidates containing GH18 domains. The Protein Disulfide Bond Enhancer (PDBE) is also his-tagged and therefore co-purifies with target proteins of interest. The DHFR plasmid, provided in the NEBExpress Cell-free E coli Protein Synthesis System, was run as a positive control for cell free expression. Visible protein bands of target sequences of interest are highlighted with a red circle.

